# Splitting sleep between the night and a daytime nap reduces homeostatic sleep pressure and enhances long-term memory

**DOI:** 10.1101/2020.09.21.301325

**Authors:** James N. Cousins, Ruth L. F. Leong, S. Azrin Jamaluddin, Alyssa S. C. Ng, Ju Lynn Ong, Michael W.L. Chee

**Affiliations:** Centre for Sleep and Cognition, Yong Loo Lin School of Medicine, National University of Singapore, Singapore; Neuroscience and Behavioral Disorders Program, Duke-NUS Medical School, Singapore; Donders Institute for Brain, Cognition & Behaviour, Radboud University Medical Centre, 6525 EN, Nijmegen,The Netherlands

## Abstract

Daytime naps have been linked with enhanced memory encoding and consolidation. It remains unclear how a daily napping schedule impacts learning throughout the day, and whether these effects are the same for well-rested and sleep restricted individuals. We compared memory in 112 adolescents who underwent two simulated school weeks containing 8 or 6.5 hour sleep opportunities each day. Sleep episodes were nocturnal or split between nocturnal sleep and a 90-min afternoon nap, creating four experimental groups: 8h-continuous, 8h-split, 6.5h-continuous and 6.5h-split. Declarative memory was assessed with picture encoding and an educationally realistic factual knowledge task. Splitting sleep significantly enhanced afternoon picture encoding and factual knowledge under both 6.5h and 8h durations. Splitting sleep also significantly reduced slow-wave activity during nocturnal sleep, suggesting lower homeostatic sleep pressure during the day. There was no negative impact of the split sleep schedule on morning performance, despite a reduction in nocturnal sleep duration. These findings suggest that naps could be incorporated into a daily sleep schedule that provides sufficient sleep and benefits learning.

## Introduction

Daytime naps consistently improve memory in a laboratory setting (Alger et al., 2012; Cellini et al., 2016; Cousins et al., 2019d; Mander et al., 2011; Ong et al., 2020; van der Helm et al., 2011), but how these benefits translate to real world sleep and learning is less well understood. The optimisation of sleep occurring before and after school classes could provide large benefits for learning (Ribeiro & Stickgold, 2014). Sleep schedules that include regular daytime naps have the potential to both alleviate learning deficits associated with poor nocturnal sleep (Cousins et al., 2018; Lo & Chee, 2020; Yoo et al., 2007), and take advantage of sleep-dependent memory benefits to the encoding and consolidation of declarative memories (Rasch & Born, 2013; Tononi & Cirelli, 2014).

Napping has been proposed as a potential remedy for insufficient nocturnal sleep (Lo & Chee, 2020), particularly during adolescence where nocturnal sleep is often below the recommended 8-10 hours on weekdays and extended on weekends in an attempt to “catch up” (Hirshkowitz et al., 2015; Yeo et al., 2019). This restricted sleep stems from a number of factors that coalesce during adolescence – including a delayed circadian cycle that is incompatible with earlier school start times (Carskadon, 2011) – and is linked to impaired cognition (Cousins et al., 2018; Cousins et al., 2019c; Lo & Chee, 2020) and academic performance (Okano et al., 2019). Unlike working adults who have limited opportunities to take an afternoon nap, 40-60% of adolescents report napping on weekdays (Calamaro et al., 2009; Jakubowski et al., 2016; Thorleifsdottir et al., 2002). This may be in response to reduced nocturnal sleep, given that more adolescents nap regularly on weekdays compared to weekends (Jakubowski et al., 2017; “National Sleep Foundation (2011) Communications technology in the bedroom,”), while those who nap regularly tend to have shorter nocturnal sleep duration (Jakubowski et al., 2017). Furthermore, experimental evidence shows that naps alleviate the negative impact of restricted sleep on several cognitive faculties (Cousins et al., 2019a; Lo et al., 2019), therefore many adolescents may already be dealing with the effects of restricted sleep by taking regular daytime naps.

While naps are primarily viewed as a consequence of deficient nocturnal sleep, or as a potential remedy for it, an alternative view is that naps could be beneficial for everyone as part of a daily schedule. There is a biological propensity to sleep in the afternoon – the circadian dip (Brooks & Lack, 2006; Monk, 2005) – and napping at this time of day is encouraged in some cultures (Ji et al., 2019). This capacity to nap is also implied by laboratory studies, where the majority of participants are capable of napping in the afternoon for 10-90 min even after a night of sufficient nocturnal sleep (Cousins et al., 2019d; Ong et al., 2020).

The barrier to understanding whether daily naps are a useful tool to optimize learning is that most prior studies examined naps in isolation, rather than integrated into a daily schedule. These studies have overwhelmingly shown that a single daytime nap enhances the consolidation of information learned before napping (Alger et al., 2012; Cellini et al., 2016; van der Helm et al., 2011), and the encoding of materials encountered after the nap (Mander et al., 2011; Ong et al., 2020). In all of these studies however, the nap forms an additional period that supplements nocturnal sleep, so that the nap condition comprises significantly greater total sleep achieved over a 24-hour period. In order to assess the benefits of daily naps as part of a regular sleep schedule, the focus must be on splitting sleep rather than supplementing sleep. That is, a fixed sleep opportunity must be split between the night and a nap, then compared to the same opportunity of purely nocturnal sleep over several consecutive days. A handful of studies investigated this with the aim of optimizing shift work in adults (Jackson et al., 2014; Kosmadopoulos et al., 2014; Mollicone et al., 2008). While none assessed learning, they did show that core cognitive functions that underscore learning such as vigilance and processing speed were unaffected by splitting sleep, and instead were determined by total sleep duration across each day.

To assess the impact of splitting sleep on learning, we recently examined performance during two typical school weeks of restricted sleep in adolescents (6.5h sleep opportunity each day), when sleep was either split between the night (5 h) and a daytime nap (1.5 h), or obtained nocturnally in one continuous sleep episode (Cousins et al., 2019b). We found that splitting sleep significantly improved declarative learning shortly after the nap in the afternoon. Importantly there were no negative effects of the split sleep schedule on performance at other times of day. These findings suggest that splitting sleep can maximize learning outcomes in those obtaining insufficient sleep, but raised the question of whether splitting sleep might also benefit long-term memory in those who are not sleep deprived.

In addition to the behavioral outcomes of splitting sleep, the impact upon sleep physiology also remains to be determined. Slow-wave sleep and spindles have been consistently linked with declarative memory consolidation (Alger et al., 2012; Ngo et al., 2013; Schabus et al., 2004) and encoding (Ong et al., 2020; van der Werf et al., 2009) in both naps and nocturnal sleep. These sleep features have been linked to synaptic downscaling in order to prevent saturation in memory networks (Tononi & Cirelli, 2014), as well as the active replay of memories to reorganize them across systems in the brain (Rasch & Born, 2013). It remains unclear how these different sleep processes are affected by splitting sleep and how they relate to long-term learning outcomes.

Addressing this, the current study replicated our prior experiment with two additional groups of adolescents (Table 1) that were afforded the minimum recommended for this age group of 8h sleep opportunity each day (8h-split and 8h-continuous), then compared their performance with groups from the prior study (6.5h-split and 6.5h-continuous) (Cousins et al., 2019b). The effect of these schedules on sleep physiology and declarative memory (picture encoding and factual knowledge task) was investigated across two simulated school weeks (Fig. 1). This included two baseline days (B_1_-B_2_), a first manipulation week (M1_1_-M1_5_) followed by 2 recovery days (R1_1_-R1_2_), then a second manipulation week (M2_1_-M2_3_) followed by recovery (R2_1_-R2_2_).

**Table 1.** Characteristics of split and continuous sleep groups.

|  | <u>Split sleep</u> |  |  |  | <u>Continuous sleep</u> |  |  |  | *P* |
| --- | --- | --- | --- | --- | --- | --- | --- | --- | --- |
|  | <u>6.5h-split</u> |  | <u>8h-split</u> |  | <u>6.5h-continuous</u> |  | <u>8h-continuous</u> |  |  |
|  | Mean | SD | Mean | SD | Mean | SD | Mean | SD |  |
| *N* | 29 | - | 24 | - | 29 | - | 29 | - | - |
| Age (years) | 16.55 | 0.74 | 16.72 | 1.16 | 16.58 | 1.12 | 16.18 | 0.88 | 0.22 |
| Gender (% male) | 51.70 | - | 45.83 | - | 51.70 | - | 48.28 | - | 0.97 |
| Body Mass Index | 20.67 | 2.80 | 20.17 | 3.13 | 21.3 | 3.5 | 20.49 | 3.08 | 0.64 |
| Daily caffeine intake (cups) | 0.55 | 0.69 | 0.63 | 0.82 | 0.58 | 0.80 | 0.72 | 1.00 | 0.87 |
| Morningness-Eveningness Questionnaire | 50.72 | 7.07 | 48.83 | 6.80 | 48.97 | 7.54 | 49.45 | 6.18 | 0.73 |
| Epworth Sleepiness Scale | 7.86 | 3.78 | 7.42 | 3.09 | 8.21 | 3.43 | 7.66 | 3.04 | 0.85 |
| Chronic Sleep Reduction Questionnaire | 36.10 | 4.66 | 37.00 | 5.23 | 35.24 | 5.96 | 36.59 | 4.04 | 0.61 |
| *Pittsburgh Sleep Quality Index* |  |  |  |  |  |  |  |  |  |
| Weekday TIB (h) | 6.78 | 0.89 | 7.36 | 1.16 | 6.85 | 1.35 | 7.59 | 1.38 | 0.33 |
| Weekend TIB (h) | 8.76 | 1.23 | 9.10 | 1.32 | 8.93 | 1.18 | 8.52 | 1.32 | 0.74 |
| Weekday TST (h) | 6.47 | 0.86 | 6.87 | 1.22 | 6.46 | 1.19 | 6.62 | 1.00 | 0.75 |
| Weekend TST (h) | 8.41 | 1.18 | 8.78 | 2.19 | 8.56 | 1.20 | 8.43 | 1.25 | 0.97 |
| Nap TST (min) | 62.93 | 65.66 | 58.54 | 60.00 | 68.52 | 64.76 | 42.59 | 53.71 | 0.42 |
| Global score | 4.17 | 1.77 | 4.22 | 1.24 | 4.48 | 1.50 | 4.48 | 1.70 | 0.82 |
| *Actigraphy* |  |  |  |  |  |  |  |  |  |
| Weekday TIB (h) | 6.84 | 1.13 | 7.22 | 0.85 | 7.00 | 0.77 | 7.21 | 0.74 | 0.32 |
| Weekend TIB (h) | 8.15 | 1.05 | 8.40 | 1.17 | 8.45 | 1.13 | 8.36 | 1.04 | 0.74 |
| Weekday TST (h) | 5.50 | 0.89 | 5.74 | 0.84 | 5.51 | 0.75 | 5.73 | 0.65 | 0.50 |
| Weekend TST (h) | 6.64 | 1.00 | 6.74 | 1.26 | 6.76 | 1.14 | 6.65 | 0.94 | 0.97 |
| Average TST (h) | 5.83 | 0.73 | 5.98 | 0.78 | 5.86 | 0.68 | 5.97 | 0.58 | 0.80 |
| SE (%) | 81.04 | 6.64 | 79.85 | 6.87 | 79.02 | 5.57 | 79.45 | 6.10 | 0.65 |

TIB = Time in bed; TST = total sleep time; SE = sleep efficiency. *P* values from one-way ANOVA and Chi-squared tests contrasting the 4 groups are listed.

**Figure 1.**
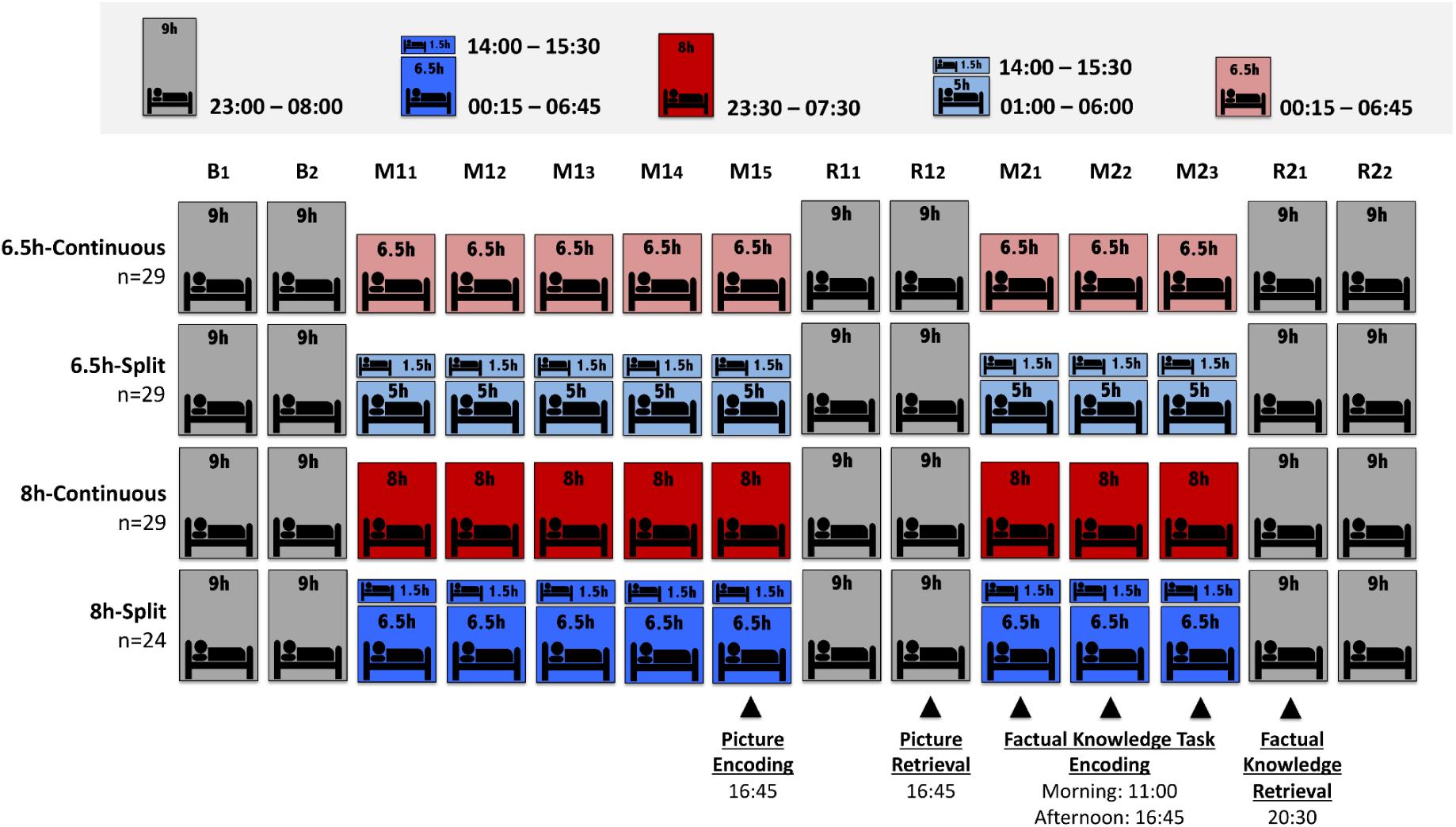
Study Protocol. All groups began the study with two baseline nights (B_1_–B_2_) of 9h nocturnal sleep opportunity. This was followed by a 5-day “school week” manipulation period (M1_1_–M1_5_) where groups diverged in terms of their sleep duration (6.5h or 8h opportunity) and distribution of sleep throughout each day (split or continuous). Continuous sleep groups slept only nocturnally, while split sleep groups obtained their sleep nocturnally with a 1.5h afternoon nap (14:00–15:30). This first week was followed by a recovery “weekend” of two days with 9h nocturnal sleep opportunities (R1_1_–R1_2_). The second “school week” was identical to the first except it lasted 3-days rather than 5-days (M2_1_–M2_3_). The study concluded with a second recovery “weekend” with 9h sleep opportunities (R2_1_–R2_2_). Participants incidentally encoded pictures at the end of the first manipulation period (M1_5_) in the afternoon and were tested on their memory after two nights of recovery (R1_2_). Participants learned factual knowledge in the morning and afternoon on each day of the second manipulation period (M2_1_–M2_3_), and were tested after one night of recovery sleep (R2_1_). Sleep physiology was assessed with polysomnography on several nights and naps throughout the protocol (B_2_, M1_1_, M1_3_, M1_5_, R1_1_, M2_1_, M2_3_, and R2_1_).

To assess memory encoding, on the fifth day of the first week (M1_5_) participants incidentally encoded pictures in the late afternoon (Fig. 2). Recognition was tested after groups obtained two recovery nights of 9h sleep opportunity. This delayed test ensured that groups were equally well rested during retrieval, and had the same post-encoding opportunity for consolidation, therefore any differences in memory could be attributed to the effects of sleep schedules specifically on memory encoding.

**Figure 2.**
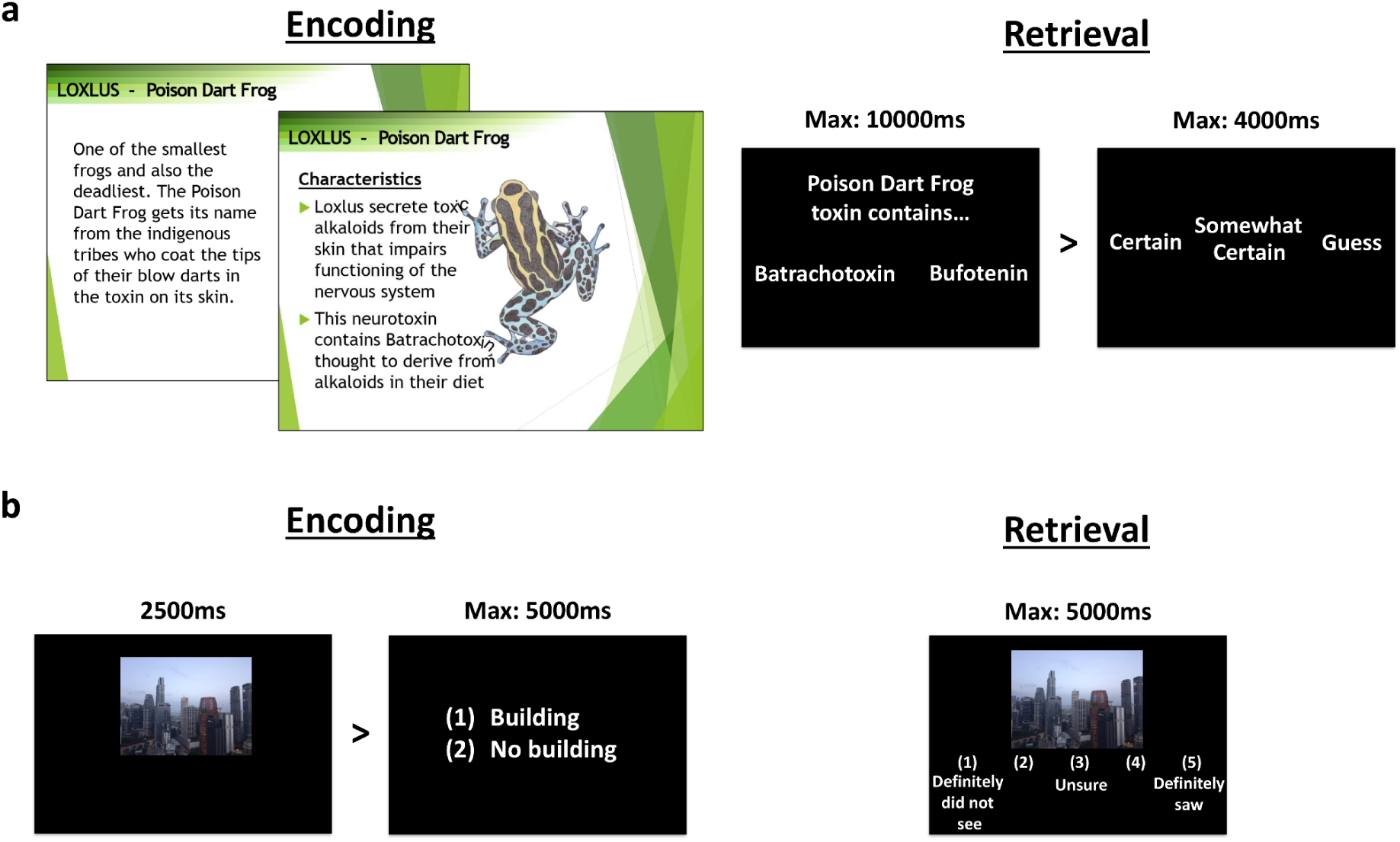
Stimuli. **a** Factual knowledge encoding involved slides with detailed information and pictures about 12 species of amphibians. Each encoding session included six species and lasted one hour. Participants were encouraged to take notes to help them learn. Retrieval consisted of 2-alternative forced choice questions of varying difficulty (360 questions in total) followed by a confidence rating. **b** For picture encoding, participants were not aware that it was a memory task and simply viewed 160 pictures, each followed by a building/no-building judgment. All 160 old images were presented again during retrieval, randomly intermixed with 80 new images. Participants were required to indicate their confidence that images were old or new on a 5-point scale.

To examine a more ecologically valid learning task, in the second week participants spent an hour learning facts about six amphibians in the morning, and an hour learning about six different amphibians in the late afternoon (Factual Knowledge Task). This was repeated across three consecutive days (M2_1_-M2_3_), then tested the following evening after one night of recovery sleep (9h sleep opportunity).

We predicted that with the increased 8h sleep opportunity, the split sleep schedule would provide a similar boost to afternoon performance as it did for both memory tasks under the restricted 6.5h sleep opportunity, evidenced by significantly better afternoon performance of the 8h-split group relative to the 8h-continuous group. We also expected that duration of sleep would impact performance, with the 8h duration groups outperforming 6.5h groups for learning in the morning and afternoon.

## Results

### Picture encoding in the afternoon was enhanced by splitting sleep, irrespective of sleep duration

The incidental encoding session occurred after 5-days in the first manipulation week on M1_5_ (16:45). Participants viewed pictures and made judgments concerning whether they contained buildings or not, for which accuracy was very high (6.5h-split= [Mean ± Standard Deviation] 0.97±0.03; 6.5h-continuous=0.94±0.06; 8h-split=0.98±0.01; 8h-continuous=0.96±0.03). A 2×2 ANOVA with sleep schedule (split/continuous) and duration (6.5h/8h) showed a significant main effect of schedule (F(1,107)=7.663, p=0.007). Splitting sleep led to significantly higher accuracy under both 8h (8h-continuous vs. 8h-split: t(51)=2.654, p=0.011, Cohen’s d (*d*)=0.732) and 6.5h sleep durations (6.5h-continuous vs. 6.5h-split: t(56)=2.230, p=0.030, *d*=0.586). There was no significant main effect of duration (F(1,107)=1.322, p=0.253) and no schedule*duration interaction (F(1,107)=0.468, p=0.495).

Our primary measure of encoding performance was picture recognition (A’) when probed at the surprise retrieval session on R1_2_ (16:45), for which no participants reported expectation of a memory test. A 2×2 ANOVA showed a significant main effect of schedule (F(1,107)=7.358, p=0.008), driven by significantly enhanced memory in split sleep groups under the 8h (8h-continuous vs. 8h-split: t(51)=2.038, p=0.047, *d*=0.562) and 6.5h durations (6.5-continuous vs. 6.5h-split: t(56)=2.456, p=0.017, *d*=0.645; Fig. 3a). There was no significant main effect of duration (F(1,107)=1.880, p=0.173), and no interaction (F(1,107)=0.341, p=0.560). These findings suggest that splitting sleep improved afternoon encoding irrespective of total sleep duration.

**Figure 3.**
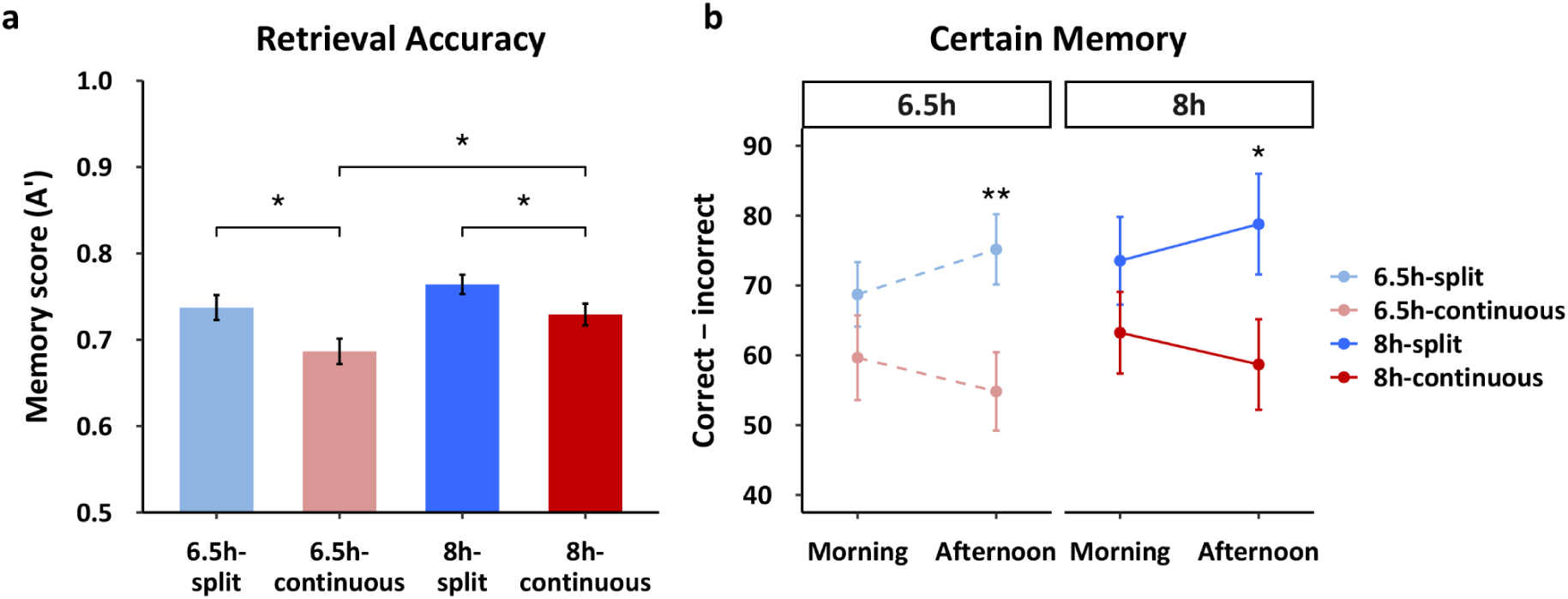
Memory performance. **a** Splitting sleep led to significantly better picture recognition (A’) for both the 6.5h and 8h durations, indicating a benefit of the nap for subsequent encoding. A longer sleep duration also provided better memory for groups that slept only nocturnally (8h-continuous vs. 6.5h-continuous), but duration did not significantly influence memory on the split sleep schedule (8h-split vs. 6.5h split). Memory performance was similar between the 8h-continuous and the 6.5h-split groups. **b** For the factual knowledge task, splitting sleep also provided significantly better memory (certain correct - incorrect) for species learned in the afternoon for both 6.5h and 8h durations. Memory for species learned in the morning did not differ between groups. Together this suggests that a split sleep schedule benefits afternoon learning with no “cost” to morning learning. Mean ± standard error of the mean (SEM). *p < 0.05, **p < 0.01

As expected, the 8h-continuous group had significantly better memory than the 6.5h-continuous group, (t(56)=2.22, p=0.03, *d=*0.581), indicating the inadequacy of 5 successive nights of sleep restricted to 6.5h. However, memory in the 6.5h-split sleep group did not differ significantly from the 8h-split (t(51)=1.425, p=0.16, *d=*0.394) or 8h-continuous groups (t(56)=0.415, p=0.68, *d=0*.109), suggesting that a 6.5h-split schedule was sufficient to return afternoon encoding performance to a form of baseline.

### Factual knowledge learning in the afternoon benefited from splitting sleep, irrespective of sleep duration

A pretest was performed shortly before learning to assess participants’ prior knowledge of the to-be-learned materials. One-way ANOVAs identified no significant group differences for pretest measures of general knowledge, specific knowledge, picture identification, subjective knowledge and subjective disgust (p>0.05). General knowledge for amphibians was relatively low (M=62.39±9.79%), but was significantly above chance (p<0.001). Specific knowledge (M=51.08±9.54%) did not differ from chance (p>0.05) while picture identification (M=53.85±14.25%) was significantly above chance (p<0.05). Subjective knowledge ratings were very low (M=1.31±0.59) with a rating of 1 representing no knowledge, while participants indicated moderate levels of disgust for the amphibian species (M=4.98±2.18). Participants also rated their subjective alertness, focus, ability and motivation during all subsequent learning sessions (Supplementary Figure S1 and Table S1).

The morning and afternoon learning sessions took place over three consecutive days (M2_1_-M2_3_). Memory was tested after one night of recovery sleep on day R2_2_ (20:30) via two-alternative forced choice questions followed by confidence ratings (certain, somewhat certain and guess). Consistent with prior work using this task (Cousins, et al., 2019b, 2019c, 2019d), our analysis focussed on certain responses that were corrected for response bias (correct-incorrect). We conducted a 2×2×2 mixed ANOVA to examine the effects of schedule (split/continuous), duration (6.5h/8h), and time (morning/afternoon) on certain memory (correct−incorrect). We found a significant main effect of schedule (F(1,107)=6.816, p=0.010), but no effects of duration (F(1,107)=0.490, p=0.486) and time (F(1,107)=0.049, p=0.825). Only the schedule*time interaction was significant (F(1,107)=10.108, p=0.002), while the 3-way and all other 2-way interactions were not (p>0.05).

We first examined the effect of the nap by comparing memory performance between split and continuous groups for each duration separately. Compared to the continuous sleep groups, split sleep groups in both the 8h and 6.5h durations had significantly better memory of the species learned in the afternoon (8h-continuous vs. 8h-split: t(51)=2.074, p=0.043, *d=0*.572; 6.5h-continuous vs. 6.5h-split: t(51)=2.700, p=0.009, *d=0*.709; Fig. 3b). In contrast, there were no significant group differences for morning learning (8h-continuous vs. 8h-split: t(51)=1.198, p=0.236, *d=0*.331; 6.5h-continuous vs. 6.5h-split: t(56)=1.189, p=0.239, *d=0*.312).

Next, we explored the difference in performance for material learned in the morning and afternoon within each group. Figure 3b shows that numerically, memory improved from morning to afternoon for the split sleep groups, while it declined for the continuous sleep groups. This change was significant for the 6.5h-split group (t(28)=2.088, p=0.046, *d=0*.388), but not the other three groups (p>0.05).

Finally, to fully characterize the nap benefit for afternoon learning we compared memory for species encoded in the afternoon across all groups. Unlike the picture encoding task, sleep duration did not have a significant effect on afternoon performance as there were no significant differences between continuous sleep groups (8h-continuous vs. 6.5h-continuous: t(56)=0.451, p=0.654, *d=0*.119) or split sleep groups (8h-split vs. 6.5h-split: t(51)=0.422, p=0.675, *d=0*.120). Remarkably, the 6.5h-split group significantly outperformed the 8h-continuous group (t(56)=2.01, p<0.05, *d=0*.527).

Overall, these findings are consistent with the picture encoding task and suggest that splitting sleep is an effective strategy to improve afternoon learning without any negative impact on morning performance.

### Splitting sleep relieved homeostatic sleep pressure, despite a reduction in sleep duration

Sleep was recorded and analysed on eight days during the protocol (Fig. 4). Splitting sleep significantly reduced total sleep time (TST) by 6-21 min each day, but in turn it also reduced the amount of homeostatic sleep pressure accumulated during the day for both sleep restricted and well-rested conditions (Fig.5 and Fig. 6)

**Figure 4.**
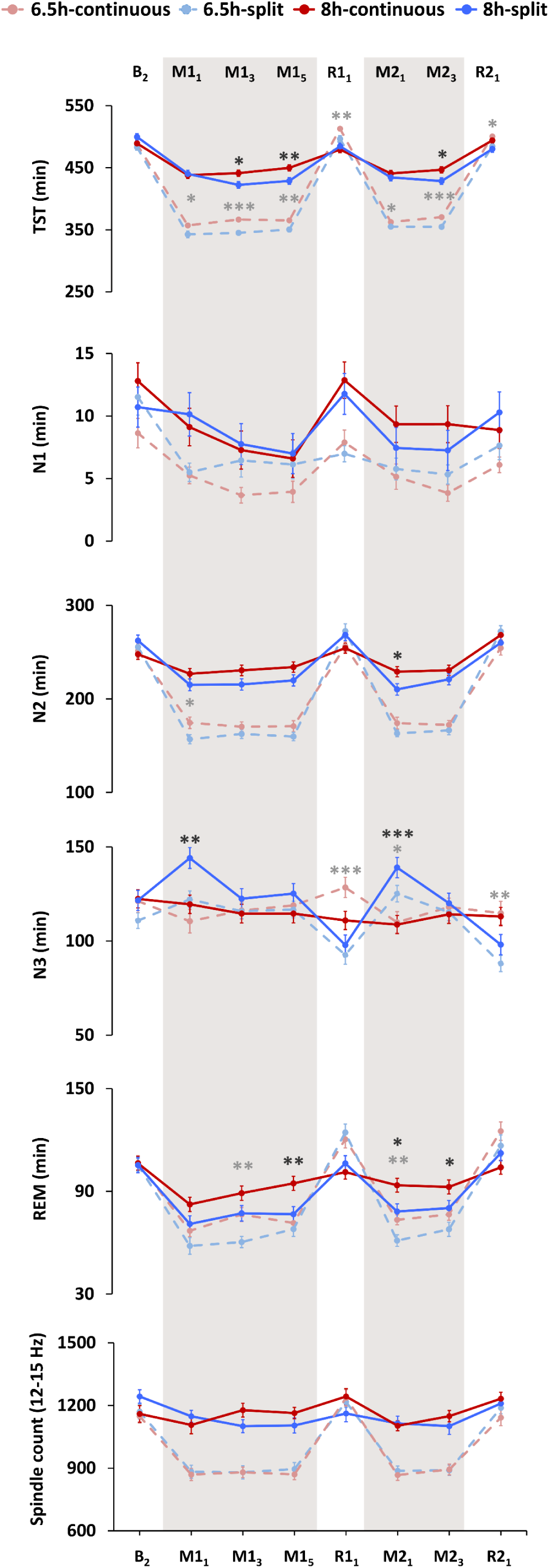
Sleep macrostructure as assessed with polysomnography. Nocturnal sleep periods were combined with the subsequent nap to compare sleep features within each 24-hour period. Splitting sleep was associated with significantly reduced total-sleep-time (TST) throughout the protocol, as well as reduced REM and N2 sleep duration on some manipulation days. Increased N3 was observed on recovery days in the 6.5h-continuous group, suggesting a greater accumulation of homeostatic sleep pressure in this group. Splitting sleep led to increased N3 at the beginning of each manipulation period. This was due to N3 during the first nocturnal sleep period being similar between groups (i.e., because this sleep period occurred before any naps had taken place), therefore the N3 achieved by split sleep groups during the subsequent nap led to overall longer duration during that 24h period. Sleep spindles were unaffected by splitting sleep. Sleep was assessed on the second baseline night (B_2_), sleep manipulation nights and naps (M1_1_, M1_3_, M1_5_, M2_1_, M2_3_; shaded areas), and recovery nights (R1_1_, R2_1_). Mean ± SEM. ***p < 0.001, **p < 0.01, *p < 0.05 for contrasts between split and continuous groups under the 6.5h duration (light grey asterisks) and the 8h duration (dark grey asterisks).

The reduction in TST was primarily due to reduced nocturnal rapid-eye-movement (REM) and N2 sleep that was not recovered during the naps. Nap sleep architecture did not differ between the two split sleep groups (6.5h-split vs 8h-split: p>0.05), with an average of 76-77 min TST, 33-34 min of N2 and 30 min of N3 sleep.

We previously showed that the accumulation of homeostatic sleep pressure could be observed in nocturnal weekday N2 latency, sleep efficiency (SE) and slow-wave energy (SWE; 0.6-4Hz), as well as weekend recovery sleep measure of nocturnal TST, SE and wake after sleep onset (WASO) (Ong et al., 2017). We assessed these 6 metrics in our groups by examining the change between baseline (B_2_) and M1_5_ (N2 latency, SE, SWE), and between B_2_ and R1_1_ (TST, SE and WASO). Together these measures indicated reduced sleep pressure in both split sleep groups (Table 2; Fig. 5).

**Table 2.**
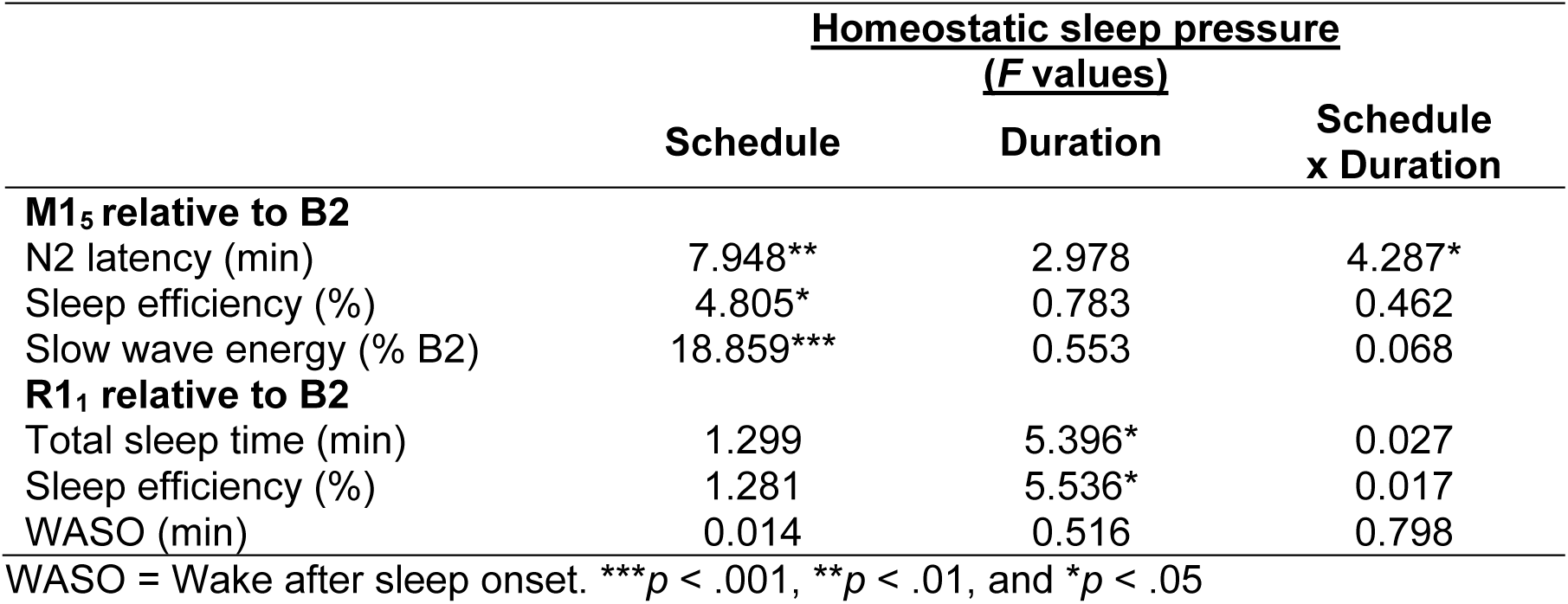
Main and interaction effects of schedule (split/continuous) and duration (6.5h/8h) on metrics of homeostatic sleep pressure.

**Figure 5.**
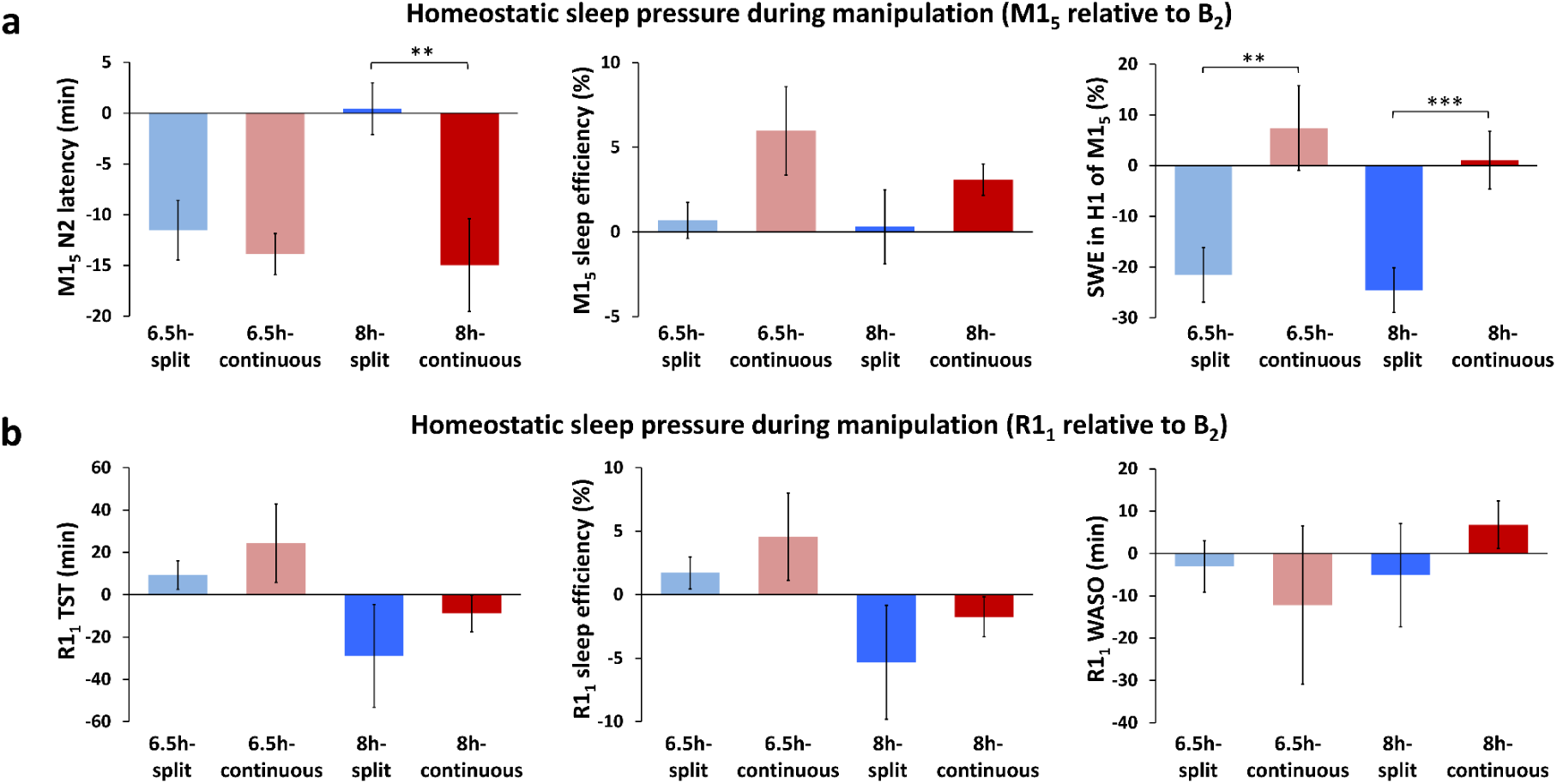
Homeostatic sleep pressure. **a** Splitting sleep led to a relative decrease in the amount of homeostatic sleep pressure by the end of the first manipulation week (M1_5_). This was evidenced by a smaller decrease from baseline (B_2_) in nocturnal N2 latency and a larger decrease in slow-wave energy in the first hour of nocturnal sleep (H1). Sleep efficiency was also significantly lower under a split sleep schedule at levels that were similar to baseline (Table 2 for main effects). **b** Splitting sleep produced numerically lower total-sleep-time (TST) and sleep efficiency during the first recovery night (R1_1_), which would suggest lower sleep pressure, but these differences were not statistically significant. There were no systematic group differences in wake-after-sleep-onset (WASO) during recovery sleep. Mean ± SEM. ***p < 0.001, **p < 0.01, *p < 0.05 for contrasts between split and continuous groups under 6.5h (light grey asterisks) and 8h durations (dark grey asterisks).

**Figure 6.**
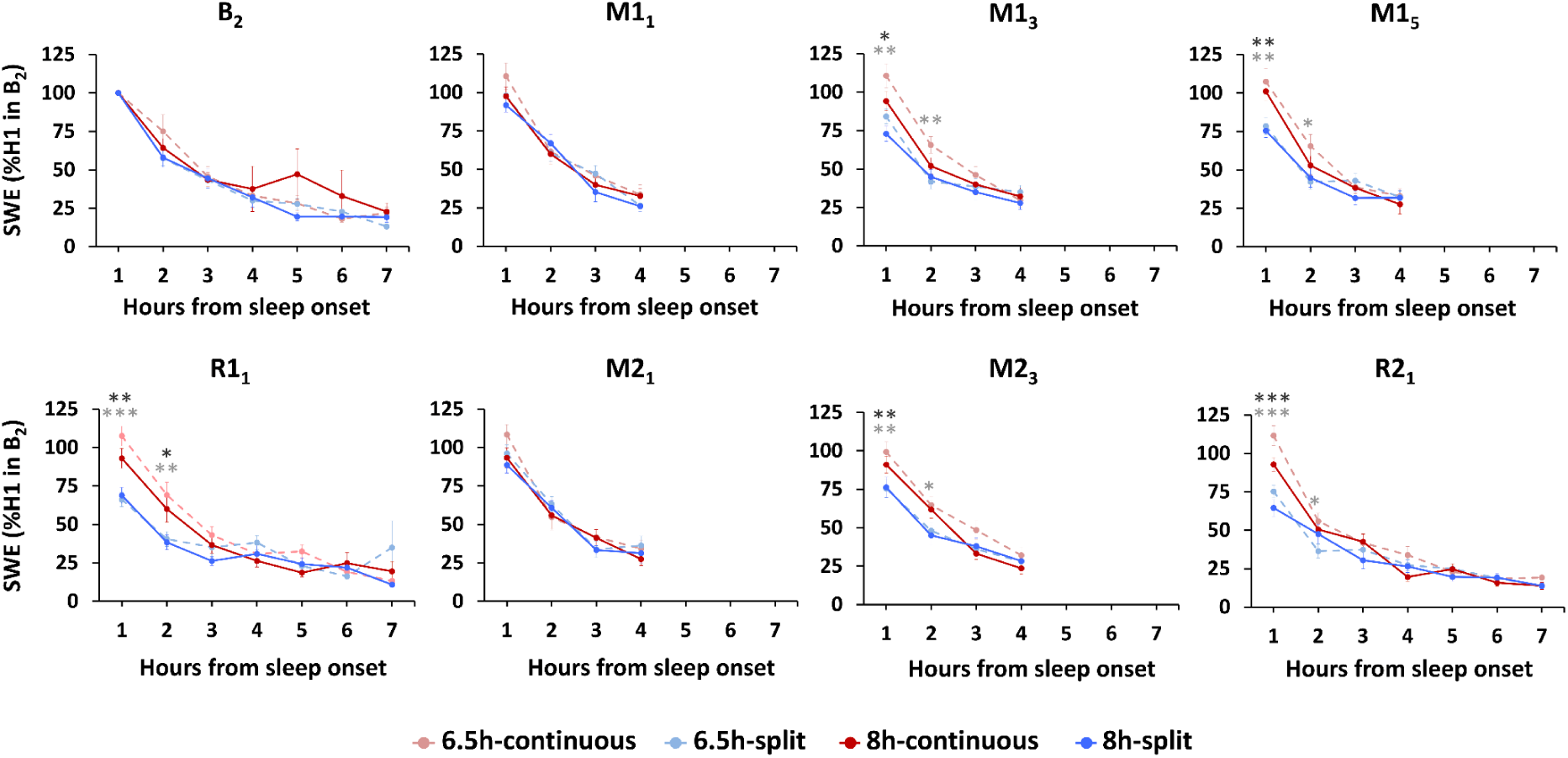
Slow wave energy (SWE) by hour from the onset of N2 sleep (normalized to SWE in the first hour [H1] of the baseline night [B2]). The split sleep schedule was associated with lower SWE in the first hour of nocturnal sleep on manipulation nights that were later in the week (M1_3_, M1_5_ and M2_3_) and recovery nights (R1_1_ and R2_1_). This suggests reduced sleep pressure for both the 6.5-split and 8h-split groups. On nights that did not have a preceding nap (M1_1_, M2_1_), there were no significant differences in SWE for the first hour of sleep (ps>0.214). Across all days and under all sleep schedules, the amount of SWE decreased as sleep progressed and levelled off after approximately 4 h of sleep. This would indicate that sleep pressure experienced by participants after waking in the morning would be similar across groups. Mean ± SEM. ***p < 0.001, **p < 0.01, *p < 0.05 for split and continuous contrasts under 6.5h (light grey asterisks) and 8h durations (dark grey asterisks).

On all nights, SWE decreased as sleep progressed and appeared to plateau after 4 h in all groups (Fig. 6). However, on the later manipulation nights (M1_3_, M1_5_ and M2_3_) and both recovery nights (R1_1_ and R2_1_), the 6.5h-continuous and 8h-continuous sleep groups showed elevated SWE compared to the 6.5h-split and 8h-split groups respectively, particularly in the first hour of sleep (ps<0.034). This suggests less accumulation of homeostatic sleep pressure in the split sleep groups over the day. By the 3^rd^ hour of sleep on all nights, groups had comparable SWE (ps>0.107).

As expected, a longer sleep opportunity led to more sleep spindles (12-15Hz; 6.5h-split vs 8h-split: ps<0.05, 6.5h-continuous vs 8h-split: ps<0.05), but splitting sleep had no effect (6.5h-split vs 6.5h-continuous: ps>0.108, 8h-split vs 8h-continuous: ps>0.162). Last, to explore associations between long-term memory and sleep, we correlated several sleep parameters with memory scores, but found no significant relationships for either memory task (p>0.05; Supplementary Tables S2 and S3).

## Discussion

We previously showed that a daytime nap enhanced afternoon learning when sleep during a typical school week was restricted to an opportunity of only 6.5 hours each day (Cousins et al., 2019b), far below the recommended daily amount of 8-10 hours for adolescents (Hirshkowitz et al., 2015). The current study examined whether that nap benefit could be applied more broadly to adolescents that are not sleep deprived, by replicating our previous study with adolescents who were afforded an adequate sleep opportunity of 8 hours each day. By combining results from both studies, we showed that splitting sleep between a nocturnal bout and a daytime nap consistently improved afternoon learning, irrespective of the total amount of sleep obtained. Crucially, this had no negative effect on participant’s capacity to learn in the morning, despite the fact that splitting sleep reduced the amount of nocturnal sleep prior to morning learning. These findings suggest that facilitating naps in education could markedly improve educational outcomes.

The nap benefit for afternoon learning was consistent not only across sleep deprived and well-rested subjects, but also over both declarative memory tasks. The factual knowledge task in particular provides evidence that the types of information encountered in daily life are more likely to be retained under a split sleep schedule. This builds upon findings that memory gains after a one hour daytime nap were equivalent to gains made from an hour of cramming (Cousins et al., 2019d), and translational research showing that regular naps taken in school can improve test scores (Cabral et al., 2018; Ji et al., 2019; Lemos et al., 2014). The ‘split sleep’ design we employed goes beyond most prior nap studies of memory (Alger et al., 2012; Cabral et al., 2018; Cousins et al., 2019d; Lemos et al., 2014; Mander et al., 2011; Ong et al., 2020) by matching the total sleep opportunity over 24 hours in those who napped (split sleep) and those who did not (continuous sleep). The memory benefits we observed can therefore be attributed to the daily naps taken under the split sleep schedule, rather than overall sleep duration as was indicated in some earlier studies of split sleep (Jackson et al., 2014; Kosmadopoulos et al., 2014; Mollicone et al., 2008). Surprisingly, memory for materials learned in the morning did not differ significantly between groups, even when comparing an 8h-continuous sleep opportunity to the 5h nocturnal opportunity of the 6.5h-split group. This suggests that morning learning is fairly resilient to these levels of sleep restriction.

The core functions of declarative memory include the initial encoding of stimuli, offline processes that strengthen and stabilise representations (consolidation), and finally the reinstatement of memories during retrieval. Research has largely focussed on sleep’s role in consolidation (Rasch & Born, 2013), but our findings most likely relate to the effect of naps on refreshing one’s capacity to encode (Mander et al., 2011; Ong et al., 2020). First, those on the split sleep schedule obtained less nocturnal sleep in which to consolidate materials learned in the afternoon, which makes it unlikely that consolidation was a contributing factor to their afternoon memory advantage. Second, all groups were afforded the same sleep opportunity during the consolidation periods after learning. For the picture encoding task in particular, this means that any group differences can be attributed to the effects of splitting sleep purely on the single encoding session that occurred at the end of the first week.

What neural mechanisms could account for this nap benefit? The synaptic homeostasis hypothesis (SHY) posits that slow-wave activity during sleep globally downscales the synaptic weights that are potentiated during encoding across the day, and this avoids saturation in memory networks (Tononi & Cirelli, 2014). It follows that encoding capacity should be maximal after waking, and subsequently declines during extended wakefulness (Cousins et al., 2018; Cousins et al., 2019c; Mander et al., 2011; Yoo et al., 2007). This framework accounts for why groups did not differ in their encoding capacity in the morning, since all achieved approximately 110-140 min of nocturnal slow-wave sleep (N3) and had been awake for a similar amount of time before learning began. This was also reflected in our analysis of slow-wave energy (SWE) where levels plateaued and groups converged after 3-4 hours of nocturnal sleep each night (Fig. 6), indicating relative normalisation of homeostatic sleep pressure across groups prior to waking in the morning. In the afternoon however, those under the split sleep schedule encoded only 1 h 15 min after waking from a nap that included approximately 30 min of N3. The nap could therefore be considered to ‘reset’ encoding capacity (Mander et al., 2011; Ong et al., 2020). Supporting this point, we found that splitting sleep significantly reduced homeostatic sleep pressure across several metrics, including reduced nocturnal N2 latency and SWE in the first hour of nocturnal sleep (Fig. 5). This suggests a reduction of sleep pressure during the daytime when memory tasks were performed. There were also hints toward these relationships in the behaviour, with parallel rates of performance decline from morning to afternoon for both continuous sleep groups. By contrast, the split sleep groups demonstrated parallel increases in performance at a similar rate (Fig. 3b). While the consistency of these changes in our four experimental groups is striking, only one of these within group changes across the day was statistically significant (6.5h-split group), therefore further studies are required to verify this variation in encoding capacity with time spent awake.

Our analysis of sleep architecture showed that splitting sleep led to a significant reduction in sleep duration over each weekday. For both sleep durations (6.5h and 8h), REM sleep was the most consistently reduced by splitting sleep, due to the fact that REM sleep occurs predominantly toward the end of nocturnal sleep and is therefore more likely to be cut by shortening sleep. Prior studies suggest that sleep duration must be reduced by many hours before cognitive deficits are observed (for review see Cousins & Fernandez, 2019), therefore the differences we observed in the range of minutes may have been too small to impact upon performance. Under a split sleep schedule, it appears that the benefits to declarative memory of reduced daytime sleep pressure outweigh any losses associated with reduced REM or N2 sleep. However, our study does not preclude possible negative effects to cognitive faculties that have been linked to these sleep stages (e.g., procedural memory (Laventure et al., 2016)) and these should be a target for future research.

The timing of the afternoon learning sessions may also be an important factor, occurring in close proximity to the circadian dip (Monk, 2005). Our findings are consistent with the idea that continuous sleep groups were impacted by this afternoon dip in performance, while the nap counteracted such effects for split sleep groups. The nap was positioned to coincide with this dip in alertness when many people have a propensity to nap (Lovato & Lack, 2010). Indeed, all of our participants could adapt to a daily napping schedule despite most reporting that they did not nap habitually in their daily lives (napping once or more per week). This capacity for all subjects to nap in the afternoon, coupled with the observed improvements to learning and memory, support the notion that napping could be an adaptive behaviour in humans.

We also found that both memory tasks displayed clear linear relationships between sleep duration and encoding performance (Fig. 3). However, this only produced a significant impairment for picture encoding, where sleep was reduced to 6.5h sleep opportunity for 5 consecutive days. Prior to this, encoding deficits had only been observed for more severe manipulations that restricted a whole night of sleep (Poh & Chee, 2017; Yoo et al., 2007) or allowed an opportunity of only 5 hours sleep for 4-5 consecutive nights (Cousins et al., 2018; Cousins et al., 2019c). Here we show a learning deficit after a relatively milder pattern of sleep loss that is common for adolescents (Yeo et al., 2019).

These findings form part of a broader picture where both sleep before and after learning contribute to long-term learning outcomes (Cousins & Fernandez, 2019), therefore both must be considered when determining the optimal sleep-learning schedules in education. Some earlier laboratory studies indicated that sleeping shortly after learning provided the best learning outcomes (Gais et al., 2006), since such memories will encounter less interference prior to sleep and therefore maximise on sleep-dependent consolidation processes (Ribeiro & Stickgold, 2014). However, when splitting sleep between the night and a daytime nap, our findings indicate that sleep occurring shortly before learning is more critical due to its beneficial effect on encoding. One caveat to consider is that we did not examine memory for materials learned in the late evening, therefore potentially these would benefit to a greater degree from consolidation processes during nocturnal sleep than the afternoon learning that we assessed.

To conclude, we show that afternoon learning is enhanced by a split sleep schedule that includes regular daytime naps, irrespective of whether individuals are sleep deprived or well-rested. Furthermore, there is no negative effect of splitting sleep on morning learning. These findings suggest that naps could be incorporated into a daily sleep schedule that provides sufficient sleep and benefits cognition. We hope this serves as a springboard for further translational research regarding the efficacy of daily naps in education.

## Methods

### Participants

Participants (n=112) comprised adolescents in the Need for Sleep (NFS) 4 (Cousins et al., 2019b; Lo et al., 2019) and NFS5 (Lo et al., 2020) studies. They were between 15-19 years old, had no history of chronic medical, psychiatric or sleep disorders, a body mass index (BMI) of ≤30kg/m^2^, were not habitually short sleepers (actigraphically assessed TIB ≥6h with weekend sleep extension ≤1h), consumed ≤5 caffeinated beverages per day, and did not travel across >2 time zones one month prior to the study.

Participants in both studies were randomised into continuous and split sleep groups with total sleep opportunity differing between the two studies (6.5 h in NFS4 and 8 h in NFS5), resulting in four groups: 6.5h-continuous (n=29), 6.5h-split (n=29), 8h-continuous (n=29) and 8h-split (n=25). Those in the continuous groups were afforded an opportunity to sleep only at night, while those in the split sleep groups had a night sleep opportunity and a 1.5-h afternoon nap. Both studies were registered clinical trials (NCT03333512 and NCT04044885).

Groups did not significantly differ in age, sex, BMI, daily caffeine consumption, morningness-eveningness preference (Morningness Eveningness Questionnaire), levels of excessive daytime sleepiness (Epworth Sleepiness Scale), symptoms of chronic sleep reduction (Chronic Sleep Reduction Questionnaire), self-reported sleep quality (Pittsburgh Sleep Quality Index) and actigraphically assessed sleep patterns (p>0.22; Table 1). Informed consent was obtained from all participants and legal guardians in compliance with a protocol approved by the National University of Singapore Institutional Review Board. Participants were compensated financially upon completion of the study.

### Procedure

The NFS4 and NFS5 study protocols tracked cognitive performance across 15 days (Fig. 1) as part of a summer camp at a local boarding school. One week prior to the protocols, participants refrained from napping and adhered to a 9h sleep schedule (23:00-08:00) confirmed with actigraphy.

Both protocols began with two baseline nights of 9h TIB followed by two cycles of sleep manipulation and recovery (9h TIB). The first cycle consisted of five nights of manipulation (M1_1_ - M1_5_) followed by two nights of recovery (R1_1_-R1_2_), while the second cycle involved three nights of manipulation (M2_1_-M2_3_) followed by two nights of recovery (R2_1_-R2_2_). On manipulation nights, the 6.5h-continuous schedule included 6.5 h nocturnal TIB (00:15-06:45), while the 6.5h-split schedule included 5 h nocturnal TIB (01:00-06:00) followed by a 1.5 h nap opportunity (14:00-15:30). The 8h-continuous schedule included 8 h nocturnal TIB (23:30-07:30) while the 8h-split schedule included 6.5 h nocturnal TIB (00:15-06:45) and a 1.5 h daytime nap opportunity (14:00-15:30). Sleep-wake patterns were monitored throughout the protocol using actigraphy. Polysomnography (PSG) was recorded on nine nights (B_1_, B_2_, M1_1_, M1_3_, M1_5_, R1_1_, M2_1_, M2_3_, and R2_1_), and daytime naps were monitored with PSG on five manipulation days (M1_1_, M1_3_, M1_5_, M2_1_, and M2_3_).

All cognitive tasks were programmed in E-Prime 2.0 (Psychology Software Tools, Inc., Sharpsburg, PA) and were administered via individual laptops in a classroom setting. Participants were required to wear earphones to minimize distraction. Timing of tasks was identical for all groups. Picture encoding took place on M1_5_, and retrieval on R1_2_, both at 16:45. Learning sessions for the factual knowledge task took place during the second week of sleep manipulation (M2_1_-M2_3_) with the morning session at 11:00 and the afternoon session at 16:45. The factual knowledge retrieval took place on R2_1_ at 20:30.

#### Picture-encoding task

The task consisted of 240 coloured scene images presented centrally on a computer screen one at a time. Half of the images depicted scenes containing buildings while the remaining half depicted landscapes with no buildings. The images were split into 3 sets of 80 images (40 buildings and 40 no-buildings). Two sets (160 images) were presented during both the encoding and retrieval sessions while the last remaining set (80 images) served as foil images to be presented during retrieval. Image sets used during encoding and retrieval sessions were counterbalanced across participants.

The encoding session took place in a single 15-min block. Participants were instructed to indicate whether or not the image contained a building. They were not informed that their memory for the images would be tested later. The images were presented for 2500ms each, followed by a response screen showing ‘(1) Building, (2) No building’. After having made a response with the corresponding keypress, an inter-trial interval of 1000ms followed before the start of a new trial. The order of presentation for the images was randomized.

The retrieval session tested the participants’ recognition of 160 ‘old’ images presented during encoding randomly intermixed with 80 ‘new’ foil images. A five-point confidence scale was presented below the images: ‘(1) Definitely did not see, (2) Probably did not see, (3) Unsure, (4) Probably saw, (5) Definitely saw’. Participants were instructed to indicate whether or not they remembered seeing the image from the previous session. Trials ended once a response was made or after a time limit of 5000ms, then followed by an inter-trial interval of 1000ms.

Responses were recorded for the encoding and retrieval sessions. Trials with incorrect responses during encoding were excluded from retrieval analyses as they indicate that images were not adequately attended to. Responses from the retrieval session were categorized into four outcome measures: (1) ‘hits’ included confidence ratings of 4 (probably saw) and 5 (definitely saw) to old images, (2) ‘false alarms’ included confidence ratings of 4 and 5 to new images, (3) ‘misses’ included ratings of 1 (definitely did not see) and 2 (probably did not see) to old images, and (4) ‘correct rejections’ included ratings of 1 and 2 to new images. To account for participants’ response bias toward old/new responses, the non-parametric signal detection measure A’ was calculated where 0.5 indicates chance performance.

#### Factual knowledge task—pretest

The pretest was administered before learning began to gauge prior knowledge of the species to-be-learned. This involved five stages: (1) Picture identification: participants identified the name of each species (two options) from an image. (2) General knowledge: 20 two-alternative forced choice questions about general characteristics of amphibians. (3) Specific knowledge: 20 two-alternative forced choice questions on to-be-learned material, similar to questions that would be administered in the final retrieval test. (4) Subjective disgust: participants rated the amount of disgust they felt toward each species on a scale of 1-9 (no disgust to extreme disgust), to control for the influence of emotion of memory. (5) Subjective knowledge: participants rated the amount of prior knowledge they had for each species on a scale of 1-9 (no knowledge to extensive knowledge). All questions were self-paced and presented in a random order.

#### Factual knowledge task—encoding

Participants learned factual information about 12 species of amphibians consisting of different animal types: three frogs (Poison Dart Frog, Flying Frog, Gray Tree Frog), three toads (Burrowing Toad, Yellow-bellied Toad, Cane Toad), three newts (Alpine Newt, Orange-bellied Newt, Great Crested Newt), and three salamanders (Giant Salamander, Green Salamander, Mud Puppy). Information regarding the characteristics of the species were adapted from their actual biology and behaviours. At the start of the session, participants were informed that all of the information they learned would be tested at a later date with example test questions shown using different species. Additionally, participants were instructed not to discuss or look up information about amphibians outside of the learning sessions.

Learning took place over three days. Each day participants underwent two learning blocks in the morning and two blocks in the afternoon, with each block focusing on a different type of animal (e.g., newts). Learning blocks lasted 30 minutes each. Participants always learned the same pairs of animal in the morning (e.g., newts and frogs) and afternoon (e.g., toads and salamanders) with the order of learning these pairs switched each day. Animal type was counterbalanced across participants for the morning and afternoon sessions.

The learning materials consisted of approximately 80 slides of factual information for each animal type presented in the form of numbered points and images. Participants were able to go through the slides at their own pace (moving forwards and backwards freely), but they were advised to keep to a certain pace to ensure that all slides were seen. To facilitate this, a timer was visible throughout the learning session with slides having markers indicating how much time should have passed in 5-min intervals.

To assist with learning, some slides asked participants to write down on a piece of paper what they could recall about the information learned in the previous slides. Participants were also encouraged to take notes which were returned to the experimenter at the end of each block. The final slide of each block instructed participants to use the remaining time to revise the information learned.

At the end of each block participants completed the Karolinska Sleepiness Scale (KSS) and were asked to rate three questions on a 7-point scale: “Was your attention focused on the task or something unrelated to the task? (1 = completely on task, 7 = completely off task)”, “How motivated were you to learn the information? (1 = completely motivated, 7 = completely unmotivated)”, and “How well do you feel you could learn the information? (1 = extremely well, 7 = extremely poorly)”. These scales measured subjective ‘focus’, ‘motivation’, and ‘ability’ respectively. Scores were subsequently inverted for analysis so that higher values represented higher levels for each measure (Supplementary Fig S1 and Table S1).

#### Factual knowledge task—retrieval

Questions were presented in a two-alternative forced choice format followed by a confidence rating (certain, somewhat certain, guess). The foil was usually the answer to the same question for a different species. Analysis focussed on certain responses because these were less likely to be contaminated with noise introduced by guessing (Cousins, et al., 2019b, 2019c, 2019d). These were corrected for response bias by subtracting incorrect from correct responses. Questions and confidence ratings remained on screen until participants gave a response or after 10 s had elapsed. In total there were 360 questions (90 questions for each animal type). Half of the questions were related to the material learned in the morning and the other half in the afternoon. Questions were presented randomly in six blocks separated by 30 s breaks. Participants were instructed to think carefully about their responses within the given time limit.

#### Polysomnography

Electroencephalography (EEG) was performed via a SOMNOtouch recorder (SOMNOmedics GmbH, Randersacker, Germany) on two channels (C3 and C4) in the international 10-20 system), referenced to contralateral mastoids. Cz and Fpz were used as common reference and ground electrodes respectively. EEG electrode impedances were kept >5 kΩ and electrooculography (EOG) and submental electromyography (EMG) impedances were kept >10 kΩ. Pulse oximetry was measured on the first night (B_1_) to assess for undiagnosed sleep apnoea.

EEG signal was sampled at 256 Hz and band-pass filtered between 0.2 and 35 Hz for EEG, and between 0.2 and 10 Hz for EOG. Automated scoring of sleep stages and artefacts were performed using Z3Score (https://z3score.com) in conjunction with the *FASST* EEG toolbox, and was also visually checked by trained staff according to standard criteria.

For each recording, the following parameters were computed: TST, N2 latency (time from lights off to N2 onset), durations of N1, N2, N3 and REM sleep, as well as sleep efficiency and WASO. As a measure of homeostatic sleep pressure, we computed slow wave energy (SWE; 0.6-4Hz) in the first hour of sleep from N2 onset by integrating power in the delta band summed across all NREM epochs, and normalized to SWE in the first hour of B_2_ (Ong et al., 2017). Automatic sleep spindle detection analysis was performed using the Wonambi Python package, v5.24 (https://wonambi-python.github.io) with an automated algorithm (Molle, Marshall, Gais, & Born, 2002). Spindle count was computed for NREM epochs using C3-A2 electrodes.

#### Statistical analysis

For picture encoding we assessed accuracy during encoding and memory at retrieval (A’) via a 2×2 ANOVA with between-subject factors of sleep schedule (split/continuous) and duration (6.5h/8h).

For the factual knowledge task, we assessed prior knowledge of the stimuli via one-way ANOVA on test scores for general knowledge, specific knowledge, picture identification, as well as subjective knowledge and disgust. Certain memory (correct-incorrect) during retrieval was analysed with a 2×2×2 mixed ANOVA including schedule (split/continuous), duration (6.5h/8h), and the within-subject factor of time (morning/afternoon). Paired and independent sample t-tests were used for follow-up comparisons. Subjective measures (alertness, focus, ability, motivation) were assessed via the same 2×2×2 mixed ANOVA.

For PSG, total-sleep-time and duration spent in sleep stages across 24 h were compared using a one-way ANOVA, and independent t-tests. Pearson’s correlations were performed between memory and sleep measures: picture encoding used the total sleep period 24 h prior to encoding (M1_5_) and recovery sleep shortly after encoding (R1_1_). Factual knowledge learning took place from M2_1_ to M2_3_, therefore we averaged the PSG-assessed sleep parameters obtained across M2_1_ and M2_3_.

All effect sizes were computed with Cohen’s d (*d*) for t-tests of key comparisons. All statistical tests were two-tailed with significance level of p<0.05.

## Acknowledgements

The authors thank Elaine van Rijn, Stijn Massar, Soon Chun Siong, June Lo, Andrew Dicom, Christina Chen, Nicholas Chee, Nicole Yu, Xin Yu Chua, TeYang Lau, Ksenia Vinogradova, Zhenghao Pu, Teck Boon Teo, Brian Teo, Jesisca Tandi, Tiffany Koa, Jessica Lee, James Teng, Kian Wong, Zaven Leow, Litali Mohapatra, Caryn Yuen, Yuvan C, Aleksi Rantanen and Karthika Muthiah for their assistance in data collection and sleep scoring. We also thank Hosein Golkashani and Shohreh Ghorbani for their help in extracting spindle metrics from polysomnography data.

