## Supplementary Materials for "Splitting sleep between the night and a daytime nap reduces homeostatic sleep pressure and enhances long-term memory"

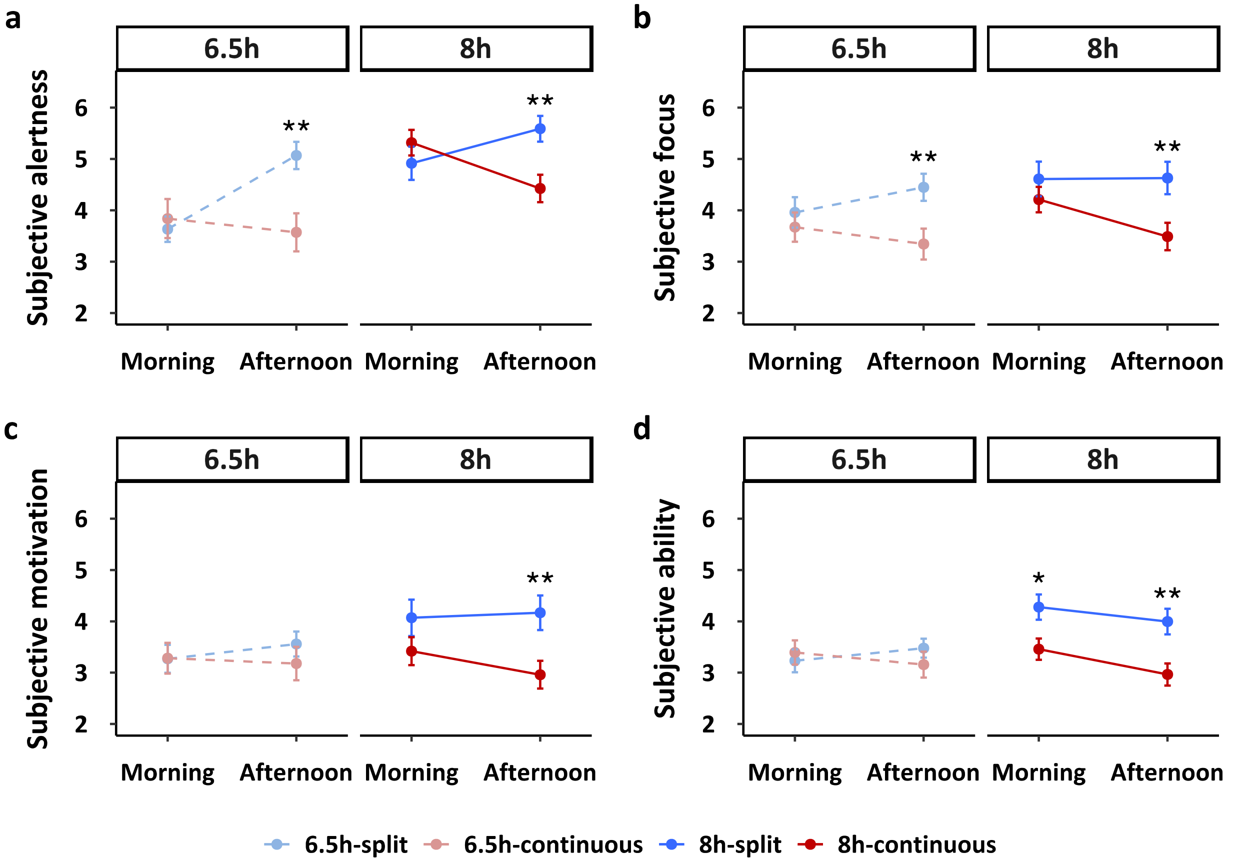


**Figure S1. Subjective measures during factual knowledge encoding.** Participants indicated their subjective alertness, focus, motivation, and learning ability during every learning session. **a** Subjective alertness was significantly higher in the afternoon under the split sleep schedule (6.5h: t(56)=-3.287, p=0.002; 8h: t(51)=-3.137, p=0.003) but similar across groups in the morning session. **b** A similar pattern emerged for subjective focus, which was higher for the split sleep groups in the afternoon (6.5h: t(56)=-2.754, p=0.008; 8h: t(51)=-2.782, p=0.008) but did not differ in the morning. **c** Subjective motivation did not differ at any point for the 6.5h groups, while it was significantly higher in the afternoon for the split sleep group under the 8h duration (t(51)=-2.824, p=0.007). **d** Lastly, subjective ability was similar for the 6.5h groups, while in contrast the 8h groups differed significantly in the morning (t(51)=-2.580, p=0.013) and afternoon (t(51)=-3.143, p=0.003). In summary, a split sleep schedule increased subjective alertness and focus in the afternoon for both the 6.5h and 8h durations, but only increased subjective motivation and learning ability in the afternoon for the 8h groups. Mean ± standard error of the mean (SEM). ***p < 0.001, **p < 0.01, *p < 0.05.

**Table S1.** Main effects and interactions for a mixed ANOVA including the factors of schedule (split/continuous), duration (6.5h/8h), and time (morning/afternoon) for participants' subjective impressions of their alertness, focus, motivation, and learning ability during factual knowledge encoding.

| | Subjective measures | | | | | | |
|---|---|---|---|---|---|---|---|
| | $F_{Schedule}$ | $F_{Duration}$ | $F_{Time}$ | $F_{Schedule\ x\ Duration}$ | $F_{Schedule\ x\ Time}$ | $F_{Duration\ x\ Time}$ | $F_{Schedule\ x\ Duration\ x\ Time}$ |
| Alertness | 2.829 | 13.790*** | 3.956* | 0.221 | 57.934*** | 9.875** | 0.100 |
| Focus | 6.664* | 1.853 | 2.869 | 0.019 | 22.729*** | 6.513* | 0.053 |
| Motivation | 3.383 | 1.216 | 0.383 | 1.710 | 8.610** | 2.823 | 0.249 |
| Learning ability | 4.680* | 2.474 | 6.770* | 3.856 | 6.591* | 7.109** | 0.882 |

****p* < .001, ***p* < .01, and **p* < .05

**Table S2:** Correlations between nap parameters and memory performance.

| | 6h-split | | 8h-split | |
|---|---|---|---|---|
| | *r* | *p* | *r* | *p* |
| **$M1_5$: Picture encoding (A')** | | | | |
| Total sleep time (min) | 0.141 | 0.474 | -0.380 | 0.100 |
| N1 (min) | -0.118 | 0.549 | 0.108 | 0.649 |
| N2 (min) | -0.025 | 0.898 | 0.218 | 0.357 |
| N3 (min) | 0.230 | 0.240 | -0.329 | 0.157 |
| Rapid eye movement sleep (min) | -0.111 | 0.575 | -0.238 | 0.313 |
| Spindle count (12-15 Hz) | -0.066 | 0.738 | 0.018 | 0.940 |
| **$M2_1$ and $M2_3$ (averaged): Factual knowledge task (certain memory)** | | | | |
| Total sleep time (min) | 0.104 | 0.591 | 0.060 | 0.790 |
| N1 (min) | 0.063 | 0.747 | -0.311 | 0.159 |
| N2 (min) | 0.106 | 0.583 | -0.162 | 0.471 |
| N3 (min) | -0.216 | 0.261 | -0.122 | 0.587 |
| Rapid eye movement sleep (min) | 0.189 | 0.325 | -0.055 | 0.807 |
| Spindle count (12-15 Hz) | 0.115 | 0.560 | -0.089 | 0.686 |

**Table S3:** Correlations between slow-wave energy (0.6-4Hz) in the first hour of nocturnal sleep (normalized to the first hour of nocturnal sleep on **$B_2$**) and memory performance.

| | **6.5h-split** | | **8h-split** | | **6.5h-continuous** | | **8h-continuous** | |
|---|---|---|---|---|---|---|---|---|
| | ***r*** | ***p*** | ***r*** | ***p*** | ***r*** | ***p*** | ***r*** | ***p*** |
| **$R1_1$:** Picture encoding (A’) | -0.369 | 0.064 | 0.227 | 0.336 | -0.079 | 0.713 | -0.349 | 0.121 |
| **$M2_1$ and $M2_3$ (averaged):** Factual knowledge task (certain memory) | 0.137 | 0.478 | 0.126 | 0.597 | -0.150 | 0.454 | -0.036 | 0.860 |
